# Unraveling axonal mechanisms of traumatic brain injury

**DOI:** 10.1101/2022.03.30.486433

**Authors:** Victorio M. Pozo Devoto, Valentina Lacovich, Monica Feole, Pratiksha Bhat, Jaroslav Chovan, Maria Čarna, Isaac G. Onyango, Neda Dragišic, Martina Sűsserová, Martin E. Barrios-Llerena, Gorazd B. Stokin

**Affiliations:** Translational Neuroscience and Ageing Program, Centre for Translational Medicine, International Clinical Research Centre, St. Anne’s University Hospital, Brno, Czech Republic; Biostatistics Department, International Clinical Research Centre, St. Anne’s University Hospital, Brno, Czech Republic; Proteomics and Mass Spectrometry Core Lab, International Clinical Research Centre, St. Anne’s University Hospital, Brno, Czech Republic; Division of Neurology, University Medical Centre, Ljubljana, Slovenia; Mayo Clinic, Rochester, MN,USA

**Keywords:** axonal swellings, traumatic brain injury, calcium, microtubules, subcortical periodic cytoskeleton, phosphoproteomics, axonal transport

## Abstract

Axonal swellings (AS) are the neuropathological hallmark of axonal injury in several disorders from trauma to neurodegeneration. Current evidence proposes a role of perturbed Ca^2+^ homeostasis in AS formation, involving impaired axonal transport and focal distension of the axons. Mechanisms of AS formation, in particular moments following injury, however, remain unknown. Here we show that AS form independently from intra-axonal Ca^2+^ changes, which are required primarily for the persistence of AS in time. We further show that the majority of axonal proteins undergoing de/phosphorylation immediately following injury belong to the cytoskeleton. This correlates with an increase in the distance of the actin/spectrin periodic rings and with microtubule tracks remodeling within AS. Observed cytoskeletal rearrangements support axonal transport without major interruptions. Our results demonstrate that the earliest axonal response to injury consists in physiological adaptations of axonal structure to preserve function rather than in immediate pathological events signaling axonal destruction.

## INTRODUCTION

Traumatic brain injury (TBI) is a major cause of disability and death (Faul and Coronado, 2015). As a consequence, the brain develops diffuse axonal injury characterized by focal enlargements within the axons known as AS (Rand and Courville, 1946; Ziogas and Koliatsos, 2018). AS are more common in unmyelinated axons (Reeves et al., 2005) and besides TBI, occur physiologically during aging (Geula et al., 2008), and in a range of pathological conditions from developmental disorders (Yagishita and Kimura, 1975) to neurodegeneration (Stokin and Goldstein, 2006).

Studies of AS report disruption (Datar et al., 2019), breakage or loss of microtubules (Tang-Schomer et al., 2010) and increased neurofilament immunoreactivity (Chen et al., 1999). Described cytoskeletal changes, coupled to observed aberrant accumulation of cargoes, suggest that AS are the aftermath of impaired axonal transport (Cross et al., 2019; Tang-Schomer et al., 2012). This hypothesis, however, is challenged by studies showing accumulation of cargoes without cytoskeletal disruption (DiLeonardi et al., 2009) and AS lacking significant accumulation of cargoes (Beirowski et al., 2010).

Since early descriptions of increased axonal Ca^2+^ levels following injury (Borgens et al., 1980), the role of Ca^2+^ in axonal injury has been the subject of intense research. Ca^2+^ surge has been observed either within AS (Barsukova et al., 2012) or across the entire axon (Staal et al., 2010; Stirling et al., 2014). Different sources have been proposed to underlie the observed Ca^2+^ increase: the extracellular milieu through mechanosensitive channels (Gu et al., 2017), voltage-gated Ca^2+^ channels (Yuen et al., 2009) and the Na^+^-Ca^2+^ exchanger (Wolf et al., 2001), and intracellularly through the ER Ca^2+^ stores (Staal et al., 2010). Studies have also shown that attenuating Ca^2+^ increase after injury reduces axonal degeneration (Ribas et al., 2017; Stirling et al., 2014). Increased axonal Ca^2+^ levels have in fact been shown to affect cytoskeletal organization through several mechanisms involving microtubules stability (Gu et al., 2017), calpain activation (Ribas et al., 2017) or calcineurin (Staal et al., 2010). Apart from Ca^2+^, inhibition of ROCK (Hemphill et al., 2011) reduced swelling formation in response to axonal injury. This finding, together with observed RhoA activation following TBI (Dubreuil et al., 2006), suggests roles of the RhoA/ROCK signaling in the AS formation.

To date, several experimental settings have been developed to study mechanisms of AS formation. Most of our current knowledge comes from *in vitro* models where neurites are stretched using dynamic substrate (Wolf et al., 2001) or bent using pressurized air (Chung et al., 2005), magnetic tweezers (Hemphill et al., 2011) or media puffing (Gu et al., 2017). However, some characteristics such as lack of discrimination between axonal or dendritic projections, or the use of neurons which are not fully mature, can be a hindrance to reproduce and elucidate the pathophysiology of TBI. Furthermore, omic approaches, apart from few exceptions (Garland et al., 2012; Nijssen et al., 2018), have not been fully exploited to study axons under normal or pathological conditions. This can be attributed mainly to the difficulty in both the isolation of an exclusive axonal fraction and the collection of sufficient material to perform a reproducible analysis. The advancement of new approaches is fundamental to understand molecular mechanisms of the very first moments of the axonal response to injury and thus to shed light to the pathology of TBI.

We here developed a human mature neuron-based system to study the immediate effects of traumatic injury on the axonal structure and function. We also generated an algorithm which allowed us to assess the real-time dynamics of the AS formation and examine the role of Ca^2+^. The system offered the unique opportunity to describe the axonal proteome, identify its phosphorylation changes and predict the kinases that are activated and deactivated in the immediate axonal response to injury. Driven by these results, we used superresolution microscopy to characterize the earliest cytoskeletal changes taking place in the axon. Finally, we assessed transport dynamics to understand the functional consequences of the traumatic injury.

## RESULTS

### Set-up of a novel axonal injury system

To investigate the immediate axonal response to injury in real-time, we coupled a microfluidic chamber (Taylor et al., 2010) harboring an exclusively axonal compartment to a syringe pump (Figure 1A). In this system, the negative pressure applied by the pump produces hydrodynamic force in the flow channel, which intersects the axons perpendicularly. To validate the system, we used fluorescent beads to measure the speed of the media in the flow channel (Figures 1B, S1A and Video S1). Pump flows between 30 and 200 µl/min produced maximum speeds similar to the ones predicted by the continuity equation and the finite element analysis. To estimate the stress to which axons are subjected, we placed urethane pillars into the flow channel and measured their displacement under the maximum pump flow of 400 µl/min (Figures 1C, S1A and Video S2). The stress reached 200 Pa, which was comparable to the values obtained theoretically using finite element analysis.

**Figure 1.**
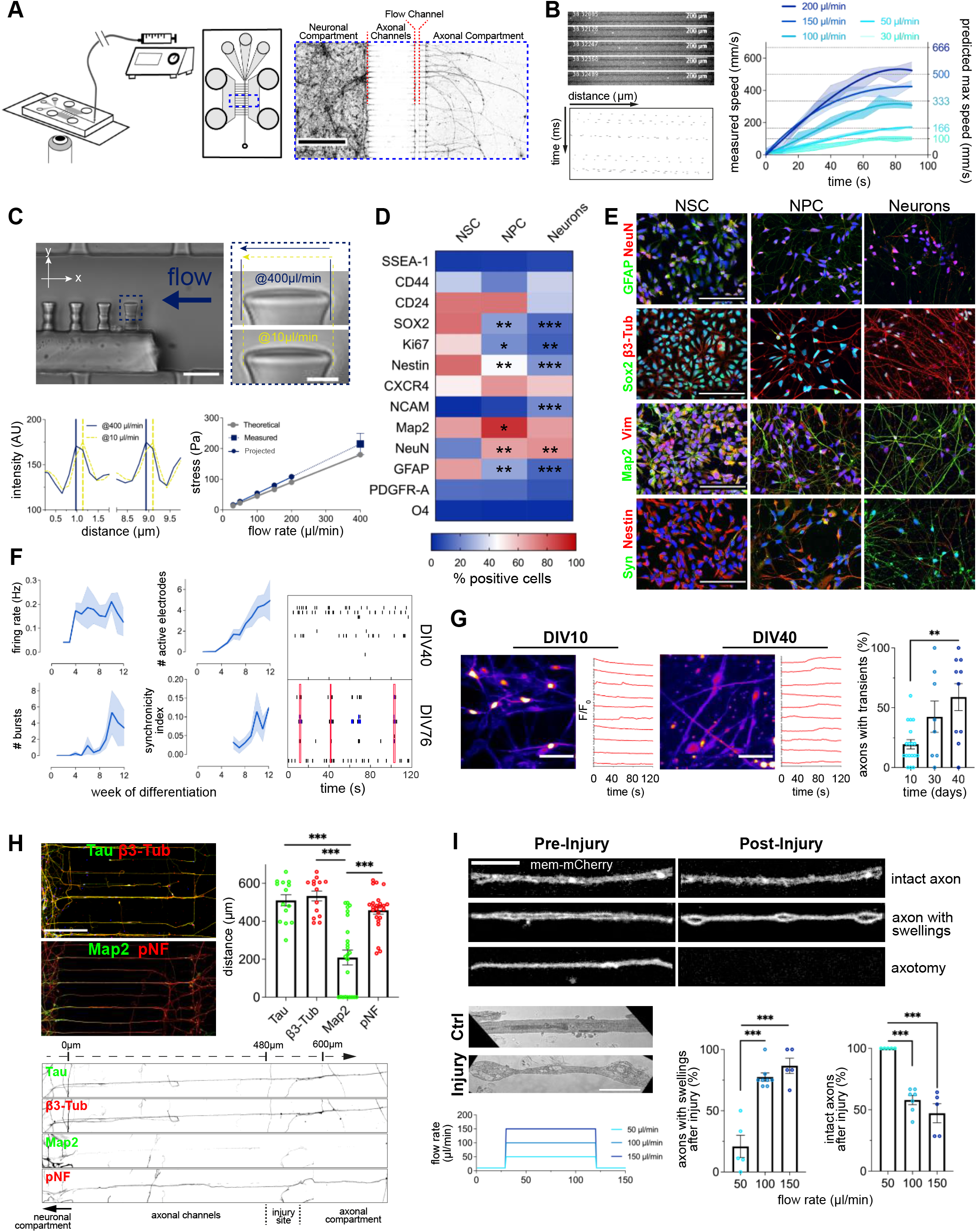
Set-up of a novel axonal injury system. (A) Schematic representation of the injury system. Image of microfluidic chamber compartments with WGA stained neurons (scale bar: 500 µm). (B) Quantification of the speed of the media at different pump flow rates by tracking fluorescent beads in the flow channel (n=3). (C) Calculation of the maximum stress applied in the flow channel by quantification of the urethane pillar bending (n=3, scale bar: 20 µm, close-up: 3 µm). (D and E) Flow cytometry quantification (D) and immunofluorescence images (E) of the expression and localization of neuronal lineage markers through the differentiation stages from NSCs to mature neurons (n=3, scale bar: 100 µm). (F) Electrophysiological activity during neuronal terminal differentiation recorded on MEA plates (n=3, 4 wells/n). (G) Quantification of Ca^2+^ transients in cultures incubated with Fluo-4 AM through terminal differentiation (n=2, 5 recordings/n, scale bar: 50 µm). (H) Immunostaining against Tau, β3-Tub, Map2 and pNF (SMI31) in the microfluidic chambers. Axonal and dendritic length measurements from the neuronal compartment to the axonal compartment (n=3, 5-10 projections/n, scale bar: 200 µm). (I) Effects of different pump flow rates on AS generation and axotomy. Axons of mem-mCherry transduced neurons imaged before and after different pump flow rates for 90 s (n=5, scale bar: 10 µm). TEM of axons in the flow channel (scale bar: 2 µm). Data are mean±SEM (*p<0.05, **p<0.01, ***p<0.001). Statistical comparisons were performed using one-way ANOVA followed by Dunnett’s multiple comparison test against NSCs group (D) or DIV10 group (G), or the Tukey’s multiple comparison test (H and I). See also Figure S1.

We next examined differentiation, enrichment and maturity of the human neurons used in our system. At the mRNA level, early stage markers of differentiation showed a gradual decrease, while markers of neuronal maturity showed increased expression across the differentiation stages (Figure S1B). At the protein level, assessment of pluripotency, mesodermal, neuroblast, neural lineage, glial and neuronal markers by FACS showed neuronal enrichment and maturity of terminally differentiated neurons further supported by the immunostaining (Figures 1D, 1E and S1C, S1D). Electrical activity was first registered during the 3^rd^ week of maturation and continued increasing throughout the 12 weeks in culture (Figure 1F), while the bursts first appeared during the 5^th^ week followed by development of interneuronal connectivity. These neurophysiological parameters were accompanied by an increase in the prevalence and intensity of the Ca^2+^ transients throughout the 40 days of neuronal differentiation (Figures 1G, S1E and Video S3).

We last tested whether projections at the site of injury correspond exclusively to the axons (Figure 1H). Immunofluorescence showed β3-tubulin, tau and phosphorylated neurofilament heavy chain, but not MAP2 immunoreactivity at the intersection and beyond the flow channel. We finally labeled cellular membrane using palmitoylated mCherry (mem-mCherry) and quantified AS and axotomy at different flow rates (Figure 1I). We identified 100 µl/min as the optimal flow rate to generate AS without significant axotomy. To confirm that mem-mCherry labeled AS indeed correspond to the actual AS, we performed TEM and observed focal enlargements of the axonal shafts as previously described (Figure 1I).

### Waxing and waning behavior of the axonal swellings

For real-time analysis of the axonal morphological changes, we developed an algorithm to track mem-mCherry or cytosolic-GFP transduced axons during the 30 s pre-injury (10 µl/min), 90 s injury (100 µl/min) and 150 s post-injury (10 µl/min) stages (Figure 2A). The algorithm enables detection of the AS by dividing axons into segments of equal length and measuring the width of each segment. A segment presents a swelling if the thickness of the shaft in that segment is at least 50% larger than the median width of all the segments (Figures 2B and 2C; Videos S4 and S5).

**Figure 2.**
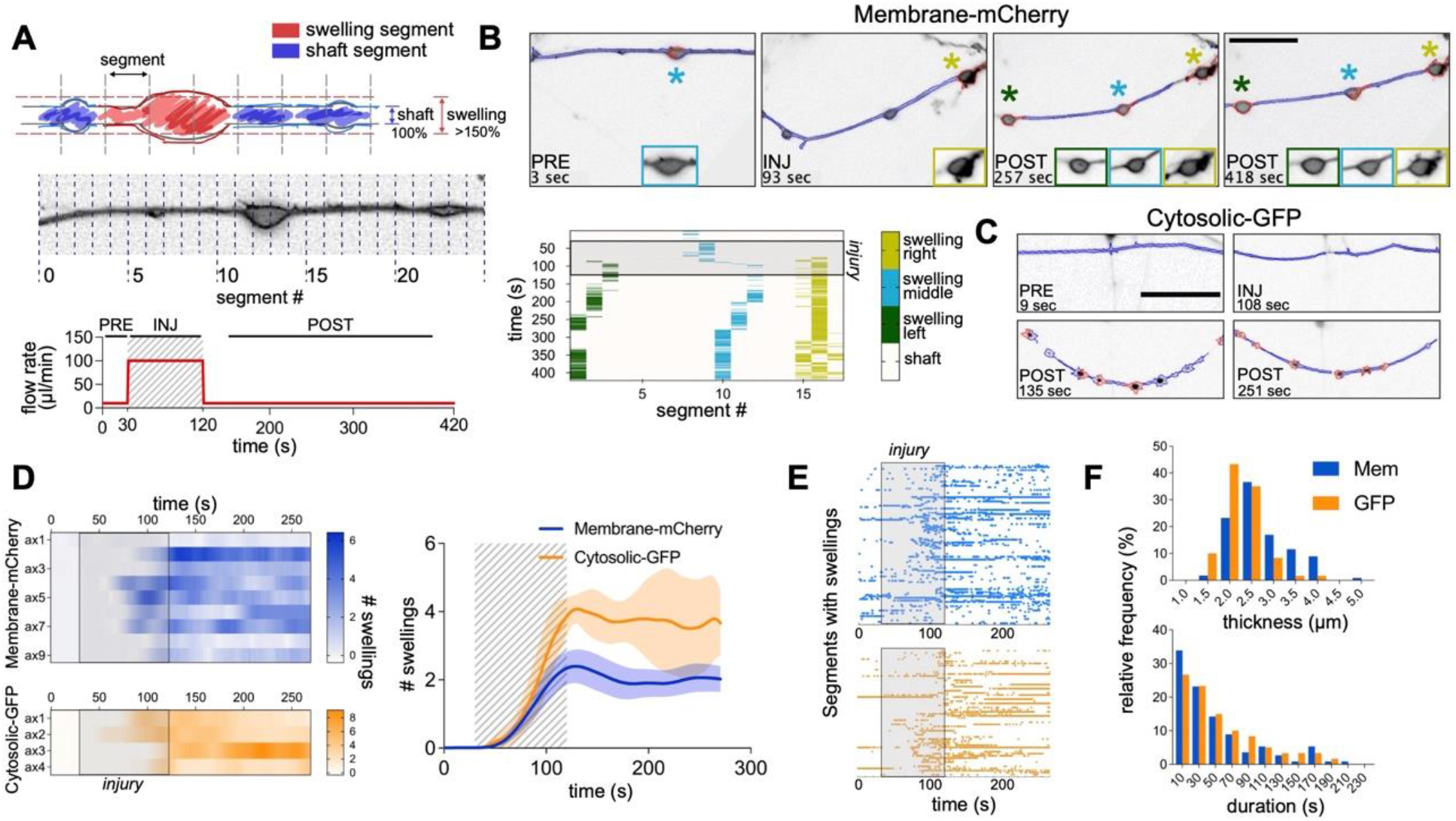
Wax and wane behavior of the axonal swellings. (A) Schematic representation of the analytical method and the injury protocol for the AS detection. The axon is divided into segments of equal sizes. For a swelling to be detected, the width of the axon in that segment has to be greater than 1.5x of the median axonal shaft width in that frame. (B) AS tracking method of axons subjected to injury. Cultures were transduced with mem-mCherry to evidence the axolemma. Matrix shows tracking of the swellings in time. Shaded area corresponds to the injury stage (from 30 to 120 s, scale bar: 20 µm). (C) Same method as in B applied to the axons transduced with cytosolic-GFP (scale bar: 20µm). (D) Quantification of the mean number of swellings per axon in time for mem-mCherry and for the cytosolic-GFP stained axons. Right plot: mean number of swellings per treatment (n=9 for mCherry, n=4 for GFP). (E) Rasterplot pooling together all the segments of the axons where swellings formed for mem-mCherry and cytoslic-GFP. (F) Frequency distribution analysis of swelling thickness and duration. Data are mean±SEM.

We observed that mem-mCherry labeled projections start developing AS approximately 50 s from the beginning of the injury and throughout the post-injury stage (Figure 2D). The same axonal behavior was observed when the neurons were transduced with cytosolic-GFP. AS developing in response to injury were most commonly short-lived, demonstrating a waxing and waning behavior (Figure 2E). Quantification of the duration of the individual AS showed that majority lasts no longer than 30 s and only a fraction persist throughout the entire post-injury stage (Figure 2F). The width of the AS ranged between 1.5-4 µm, with the most frequent widths of 2.5 µm and 2 µm for mem-mCherry and cytosolic-GFP labeled AS, respectively. In conclusion, through the use of independent markers of axonal membrane and cytosol, we demonstrate that AS that develop as a result of the hydrodynamic force, exhibited a waxing and waning behavior, with a minority persisting throughout the post-injury stage.

### Increased axonal calcium levels sustain axonal swellings

Several studies reported changes in different ion concentrations in the axon after injury (Staal et al., 2010; Yuen et al., 2009), however, it remains unsettled whether these changes are the result of membrane mechanoporation. To investigate this, we incubated intact axons and axons subjected to injury with fluorophore tagged 3, 10 and 40 kDa large dextrans (Figure S2B). No significant differences were found in the uptake of dextrans in intact versus injured axons, suggesting that the applied hydrodynamic force did not produce mechanoporation. We then asked whether injury causes any immediate changes in the axonal ion concentrations. To determine axonal Na^+^, K^+^ and Ca^2+^ concentrations in response to injury, we incubated neurons with membrane permeable SBFI, PBFI and Fura-2 ratiometric probes, and measured the 340/380 nm excitation ratios (Figures 3A and S2A). Measurements showed a significant increase in the axonal Ca^2+^, but not Na^+^ or K^+^ concentrations in response to the hydrodynamic force.

**Figure 3.**
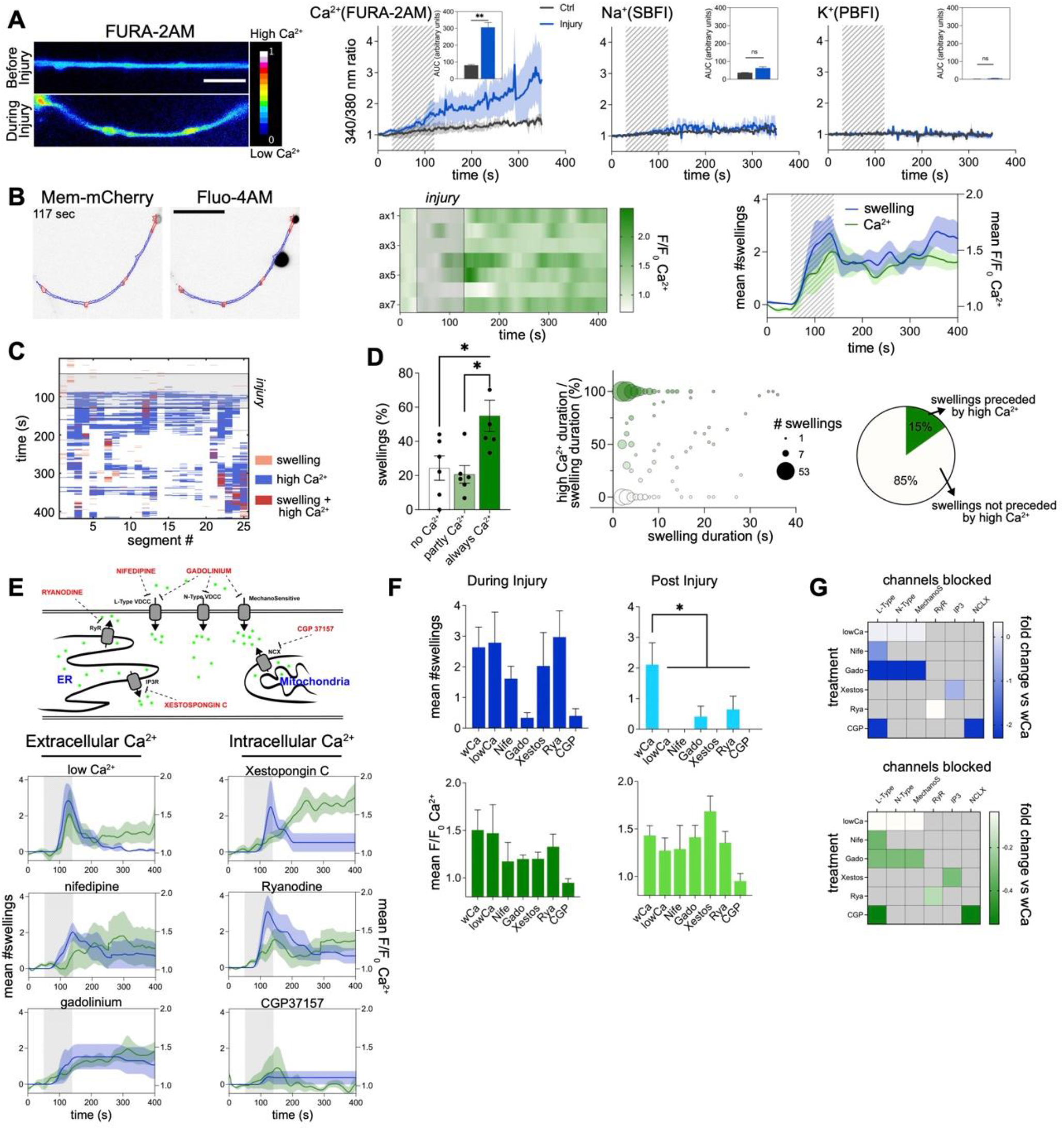
Axonal Ca^2+^ increase is required to sustain swellings. (A) Axonal Ca^2+^, Na^+^ and K^+^ levels during injury measured with the ratiometric probes FURA-2AM, SBFI and PBFI (n=4-6, scale bar: 10µm). (B) AS and axonal Ca^2+^ levels during injury. Mem-mCherry transduced neurons were incubated with Ca^2+^ sensor Fluo-4 AM. Middle: mean axonal Ca^2+^ levels during injury. Right: Mean number of swellings and Ca^2+^ levels during injury (n=7, scale bar: 20 µm). (C) AS and Ca^2+^ tracking across the axonal shaft through the injury. Example of a matrix showing detection of swellings (>150% width) and high Ca^2+^ (>1.25x) for each segment of the axonal shaft in time. (D) Percentage of AS per axon that during their duration present always Ca^2+^, part time Ca^2+^ or no Ca^2+^ (left, n=6, 10-150 swellings/n). Bubble plot shows the relationship between the duration of the AS and the presence of high Ca^2+^ (middle, n=342 swellings). Pie chart showing the percentage of AS that are preceded (2 s window) by high Ca^2+^ (right, n=342 swellings). (E) Schematic representation of the compounds used to block different sources of Ca^2+^. Axons were incubated with compounds, subjected to injury and AS formation and Ca^2+^ levels assessed (n=4-7). (F) Comparison of the effects of different compounds on the mean number of AS and Ca^2+^ intensity levels during and after injury (n=4-7). (G) Diagram showing the targets of each compound and the mean differences in AS numbers or Ca^2+^ intensity levels compared to the control levels. Data are mean±SEM (*p<0.05, **p<0.01). Statistical comparisons were performed using student t-test (A), one-way ANOVA followed by the Tukey’s multiple comparison test (D) or Dunnett’s multiple comparison test against the control group (F). See also Figure S2.

To establish dynamics of the increased axonal Ca^2+^ concentrations, we incubated neuronal cultures with mem-mCherry to visualize AS formation and the Ca^2+^ probe Fluo-4AM to monitor Ca^2+^ levels (Video S6). We observed a significant surge in the axonal Ca^2+^ concentrations during injury as well as post-injury (Figure 3B). Since this increase temporally corresponded to the AS formation, we asked if regions of the axons giving rise to AS show specific changes in Ca^2+^ levels. We found that only 54% of the AS displayed increased Ca^2+^ levels during their entire duration, 18% of the AS showed higher Ca^2+^ levels during part of their duration, while 28% showed no Ca^2+^ changes during their lifetime (Figures 3C and 3D). There was also no correlation between local Ca^2+^ presence and the AS duration, with only 15% of the AS preceded (2 s prior) by increased Ca^2+^ levels. These results show that increased axonal Ca^2+^ concentrations coincide with the AS formation, but the sites of increased Ca^2+^ levels do not always correspond to the sites of AS formation.

To investigate individual Ca^2+^ sources in the AS formation, we pharmacologically blocked several Ca^2+^ channels or transporters. Extracellular Ca^2+^ was blocked either by incubating axons with nifedipine, gadolinium or by lowering Ca^2+^ in the media, while the intracellular Ca^2+^ sources were blocked using ryanodine, xestospongin C or CGP37157 (Figures 3E, 3F, 3G and S2C). The analysis showed that none of the conditions affected AS formation during the injury, but all of them significantly decreased the number of AS in the post-injury stage. In summary, these results suggest that (i) generation of AS during the injury occurs independently from elevated axonal Ca^2+^ levels, while (ii) increased axonal Ca^2+^ levels are required for the maintenance of the AS after the injury.

### Proteomic profile of the axons following injury

To investigate the mechanisms underlying AS formation, including Ca^2+^-dependent signaling involved in their maintenance, we examined the axonal proteome and phosphoproteome in response to injury. We collected axonal fraction, composed of axons at the site and distal to the injury, and compared it to the neuronal fraction composed of axons, dendrites and somas. Considering remarkably little is known about the axonal proteome, we first performed label free quantitative mass spectrometry with the discovery (Figure 4 and Table S1) and then a confirmatory experiment comparing control (intact) versus injured axonal and neuronal fractions (Figure S3 and Table S1). The principal component analysis (PCA) and the hierarchically clustered heatmap readily distinguished axonal from the neuronal fraction, however, disclosed no injury induced changes (Figures 4A, S3A and S3B). This was further confirmed by volcano plots comparing the control to the injured axonal and neuronal compartments, which lacked reproducible quantitative protein changes between the discovery and confirmatory experiments (Figures 4B and S3C).

**Figure 4.**
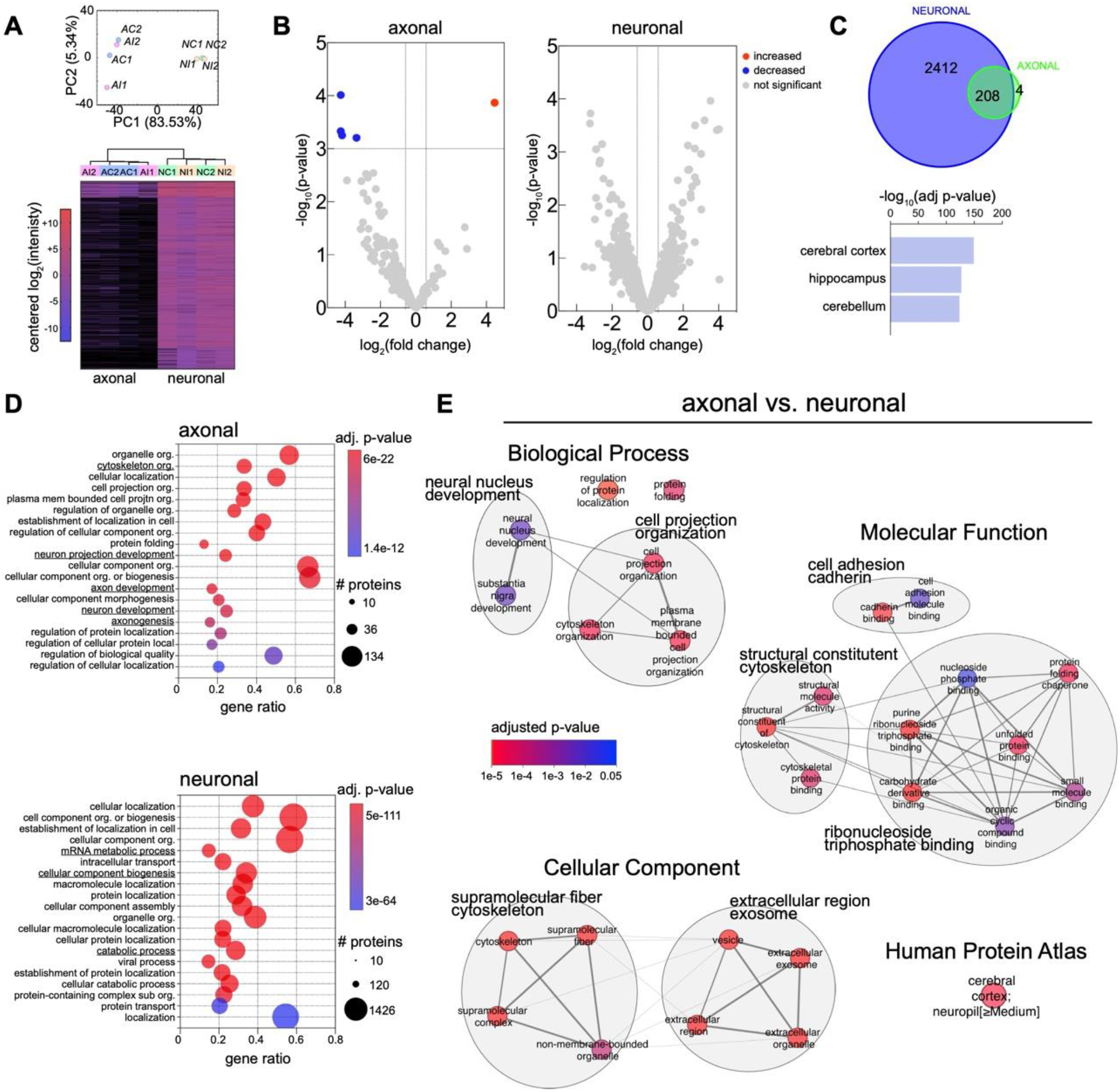
Axonal proteomic profiling before and immediately after injury. (A) Axonal and neuronal compartment fractions (A/N) in both control and injury treatments (C/I) were processed for LFQ mass spectrometry. PCA and heatmap of differentially expressed proteins (n=2). (B) Volcano plots depicting the quantitative protein changes following injury compared with control treatment for the axonal and the neuronal fractions. (C) Total number of proteins present in the neuronal and axonal fractions (upper panel). Human Protein Atlas top terms of the enrichment analysis for the neuronal fraction proteins (lower panel). (D) Top GO Biological Processes terms for the enrichment of the axonal (upper panel) or the neuronal fraction (lower panel). (E) Enrichment analysis of the axonal fraction proteins using as background the neuronal fraction proteins. Significant terms for each GO category were clustered by similarity and most representative words of the cluster’s terms assigned. See also Figure S3.

To establish the axonal proteome, we next pooled together control and injured fractions and compared the axonal to the neuronal compartment. The axonal proteome consisted of approximately 10% of the proteins identified in the neuronal fraction. Enrichment analysis showed that the three most significant terms corresponding to the neuronal fraction were related to the central nervous system: cerebral cortex, hippocampus and cerebellum (Figures 4C and S3D). Gene ontology overrepresentation analysis identified several biological processes including axon development and cytoskeleton organization enriched exclusively in the axonal fraction (Figure 4D). In contrast, mRNA metabolic process, cellular component biogenesis or catabolic process were found exclusively in the neuronal fraction. Enrichment analysis of the proteins in the axonal fraction, using the neuronal fraction as background (Figures 4E and S3E), identified clusters of terms related to cell projection organization, cell adhesion, structural constituent of cytoskeleton, ribonucleoside binding and extracellular exosome (Figures 4E and S3E). We here built an atlas cataloguing the most abundant proteins found in the axons derived from human neurons, which exhibit a proteomic profile reminiscent of the human central nervous system.

### Axonal phosphoproteome following injury

Protein phosphorylation is the major mechanism through which protein function is regulated in response to stimuli. To investigate the immediate response of axonal proteins to injury, we enriched samples for phosphopeptides and examined their phosphorylation status using mass spectrometry (Figure 5 and Table S2). In the axonal and neuronal fractions, respectively, we identified 78 and 80 differentially up-regulated phosphopeptides following injury, while 73 and 292 phosphopeptides were down-regulated (Figures 5A and 5B). We estimated that out of a total of 3345 and 3444 phosphopeptides corresponding to 1191 and 1206 proteins in the axonal and neuronal fractions, approximately 10% of the axonal and 21% of the neuronal fraction proteins underwent changes in their phosphorylation status in response to injury (Figure 5C). Enrichment analysis of the proteins that changed their phosphorylation status after injury versus all of the identified phospho-proteins showed that pathways related to the axon development are overrepresented in the axonal fraction, while no significant enrichment was found in the neuronal fraction (Figure 5D).

**Figure 5.**
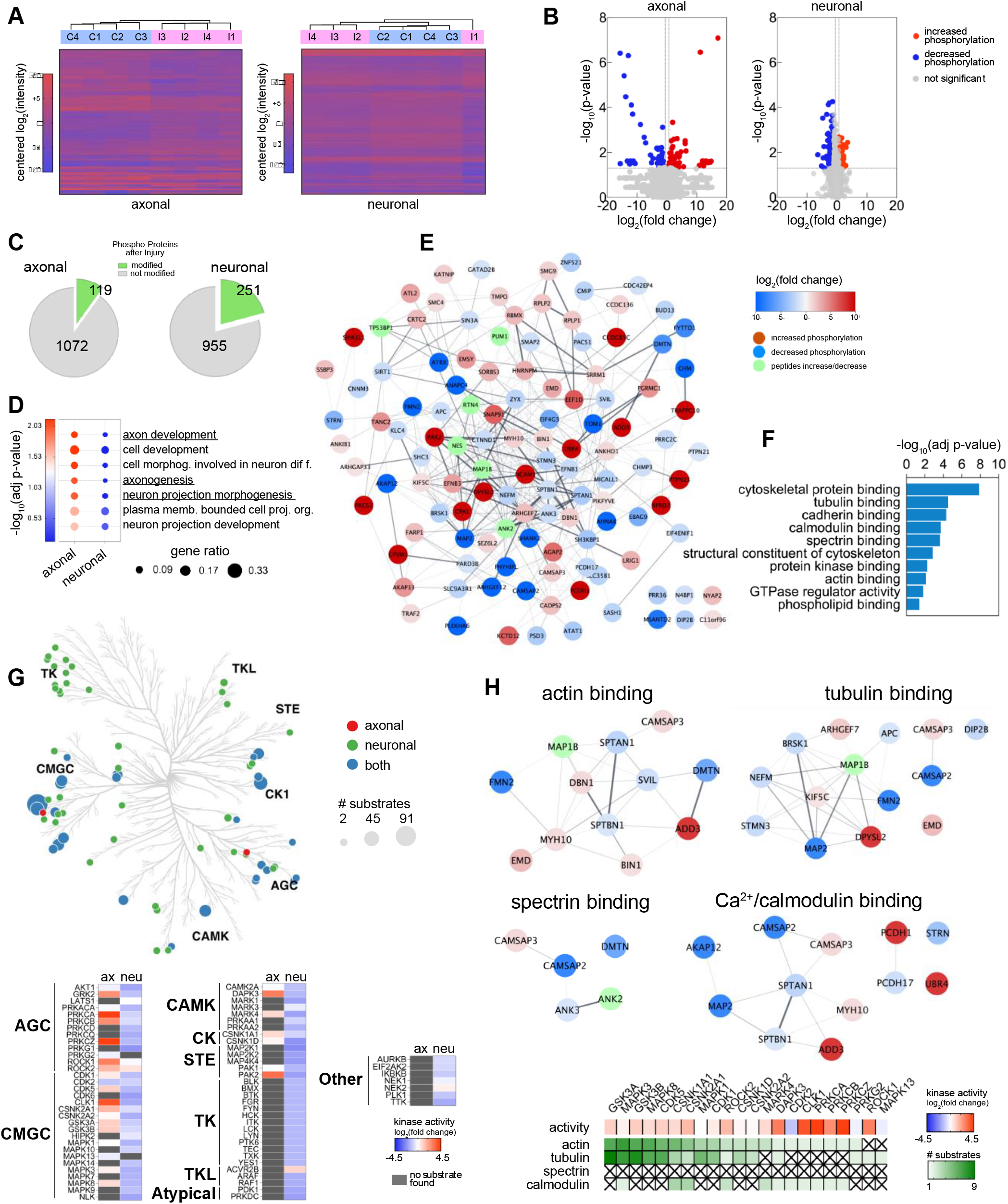
Phosphorylation changes of the axonal proteins following injury. (A) Axonal and neuronal compartment fractions (A/N) in both control and injury treatments (C/I) were enriched for phosphorylation and processed for quantitative mass spectrometry. Heatmaps of phospho-peptides levels comparing injury versus control treatment for both axonal and neuronal fractions (n=4). (B) Volcano plots of phosphopeptides levels comparing injury versus control treatment for both the axonal and neuronal fractions. (C) Pie charts depict the number of identified proteins that significantly increase or decrease their phosphorylation and the number of proteins that are not modified after injury. (D) Enrichment analysis (GO Biological Processes) of the proteins that are significantly regulated by phosphorylation after injury, using as background all the detected phosphoproteins. Comparison of significant terms between the axonal and neuronal fractions. (E) Stringplot showing all the phosphoproteins in the axonal fraction significantly changed after injury. (F) Overrepresentation analysis of the proteins in (E) showing top significant terms for GO molecular functions. (G) Kinome tree highlighting predicted kinases in both fractions, based on the significantly changed phosphopeptides as substrates (left). List of kinases and their families with the predicted activity based on their substrate’s fold change. (H) Stringplots of phosphoproteins present in specific cytoskeletal terms found in (F) and kinases dependent on the Ca^2+^/calmodulin signaling.

We then performed a detailed analysis of all the protein’s phosphorylation sites and their previously described functions (Figure 5E and Table S2). We identified phosphorylation changes in protein sites previously reported to regulate protein function and cytoskeletal organization in DMTN (S92), MYH10 (S1952), SLC9A3R1 (S280) and STMN3 (S60) (Vugmeyster and McKnight, 2009; Juanes-Garcia et al., 2015; Sun et al., 2013; Devaux et al., 2012). An enrichment analysis of Molecular Function showed that the set of proteins regulated by phosphorylation in the axon immediately after injury is overrepresented in functions such as cytoskeletal protein binding (including actin, spectrin and tubulin), binding to calmodulin and protein kinases, and regulators of GTPase activity (Figure 5F). Interestingly, no terms related to stress or degeneration were significantly enriched.

### Axonal kinome following injury

Identification of protein kinase binding among the most significantly changed molecular functions prompted us to predict putative kinases involved in the phosphoprotein changes triggered by injury (Figure 5G and Table S3). The predicted axonal kinome resulting from injury revealed that the most represented families of kinases were the CMGC (CDK, MAPK, GSK and CLK), AGC (PKA, PKG and PKC) and CAMK. The majority of the kinases found in the axonal fraction are predicted to be upregulated, with stronger activities present in PRKCA/Z/B, CLK1, PAK2, DAPK3, ROCK1/2 and GRK2. In contrast, the neuronal fraction showed a general downregulation of activity, presenting also members of the TK and TKL families of kinases.

We last linked axonal actin, tubulin, spectrin and calmodulin binding networks with their predicted kinases into discrete functional clusters (Figure 5H). We found changes in proteins participating in actin bundle stabilization or assembly (DMTN, FMN2) and linkage of actin to membrane (SVIL), proteins that have microtubule destabilizing activity (STMN3) or acetylation of tubulin (DIP2B), and proteins that participate in the localization of channels in the axolemma (ANK2 and 3). Among the calmodulin regulated proteins, CAMSAP2 and 3 bind the minus-end of microtubules and spectrin, SPTBN1 and SPTAN1 are part of the spectrin-actin subaxolemmal periodic cytoskeleton, ADD3 caps actin filaments and promotes the formation of the spectrin-actin network. In terms of kinases, we predicted their activity and the number of targeted substrates. We identified PRKCA/B/Z, ROCK1, CLK1 and DAPK3 as the overall most activated kinases. MAPK3 (ERK1), MAPK8 (JNK1), GSK3A/3B and CDK5, on the other hand, were found to target most substrates. There were no predicted kinases within the spectrin binding network.

### Reorganization of the axonal cytoskeleton following injury

Most of the phosphorylation changes of the axonal proteins following injury indicate cytoskeletal regulation. This led us to investigate the morphology and structure of the AS. We first compared morphology of the AS in control versus injured axons using scanning electron microscopy (Figures 6A and 6B). We found significant increase in the number of AS in the injured compared with control axons. These were most commonly observed as strings of up to 6 equally spaced AS with shorter inter-swelling distances. Although larger AS were occasionally observed, the majority of AS forming immediately following injury was indistinguishable in shape and size from those found in control axons (Figure S4A).

**Figure 6.**
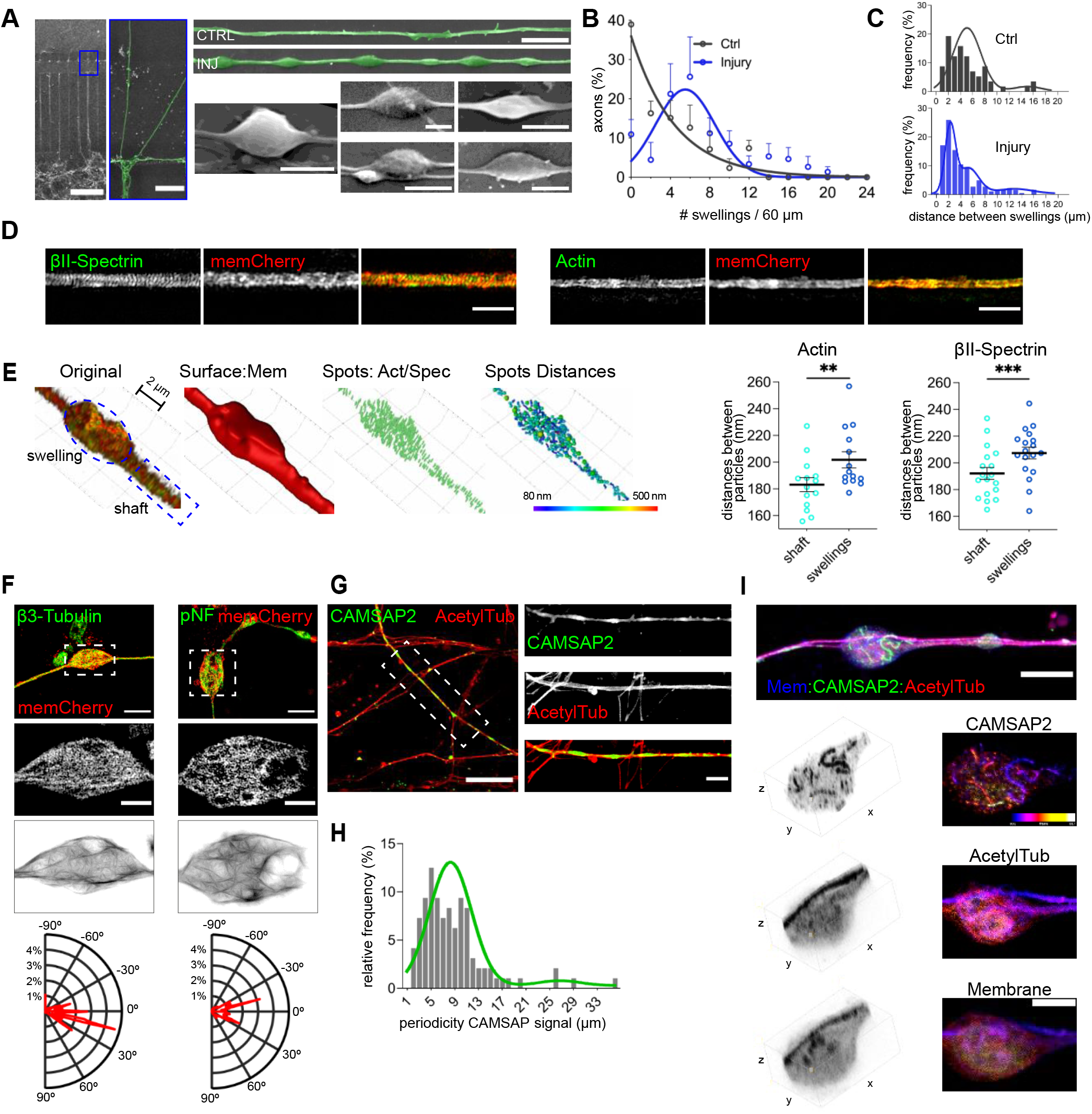
Structural reorganization of the axonal cytoskeleton following injury. (A) Representative scanning electron microscopy images of AS in control and axons subjected to injury (scale bar chamber: 200 µm, close-up: 20 µm, right horizontal: 5 µm, swellings: 2 µm). (B) Frequency distribution of the number of AS measured per axon (n=5). (C) Frequency distribution of inter-swelling distances for control and injured axons. The distribution was fitted with three Gaussian curves with means of 3.09 µm, 6.12 µm and 13.87 µm (variances: 0.75, 3.02, 8,76 µm, respectively, control n=57, injury n=181). (D) SIM images of axons transduced with mem-mCherry and immunostained for βII-Spectrin or actin (scale bar: 2 µm). (E) Identification of AS and shaft ROIs, masking for surface (mem-mCherry), identification of actin or βII-Spectrin particles and measurements of the distances between the closest neighbours. Distances in shaft vs distances in swellings for actin and βII-Spectrin (n>14). (F) SIM images of axons transduced with mem-mCherry and immunostained for βIII-tubulin or pNF. Images were processed and magnitude of directionality measured and plotted in polar plots (scale bar upper: 5 µm, middle: 2 µm). (G) Representative image of the axons immunostained for CAMSAP2 and acetylated tubulin after injury (scale bar: 20 µm, close-up: 5 µm). (H) Frequency distribution of the periodicity of CAMSAP2 staining along the axons after injury (n=96). (I) Representative image (z-projection) of an axon transduced with mem-mCherry and immunostained for CAMSAP2 and acetylated tubulin (scale bar: 10 µm, close-up: 5 µm). Data show mean±SEM (**p<0.01, ***p<0.001). Statistical comparisons were performed using the student t-test (E). See also Figure S4.

We next examined cytoskeletal structures within AS. We transduced axons with mem-mCherry, stained for actin or βII-spectrin and then imaged axons using Structured Illumination Microscopy (Figure 6D). To calculate the distance between particles, we measured the closest neighbour distance for all the particles detected inside the axon (Figures 6E and S4B). Analysis in the axonal shaft adjacent to the AS revealed 183.1 ±5.22 nm actin and 192.1 ±4.43 nm βII-spectrin periodicities consistent with previous reports (Xu et al., 2013). In contrast, actin and βII-spectrin periodicities within the AS amounted to 201.8 ±6.1 nm and 207.4 ±4.46 nm, respectively (Figures 6E, S4C and S4D). Increased distances between actin and βII-spectrin particles suggest stretching of the actin-spectrin rings in the AS. Measurements of directionality of microtubule or neurofilament tracks by polar plots showed either a linear or a criss-cross pattern within the AS (Figure 6F).

Stretched actin-spectrin rings and reorganized microtubules support a role of cytoskeletal regulatory proteins in the AS formation. To test this, we investigated CAMSAP2, which showed decreased phosphorylation after injury and was previously reported to interact with both spectrin and microtubules. Imaging revealed discrete stetches of CAMSAP2 immunoreactivity along the axons (Figure 6G). These linear stretches of CAMSAP2 adopted a snake-like shape in a subset of AS (Figure 6I). To investigate this change further, we performed real-time imaging of CAMSAP2-GFP and found bending of CAMSAP2 stretches during injury which acquired and maintained the snake-like shape after injury (Figure S4E and Video S7). These findings suggest involvement of CAMSAP2 in the microtubule reorganization within the AS, which is further supported by the periodical accumulation of CAMSAP2, reminiscent of the AS distribution along the axons (Figure 6H). These experiments demonstrate reorganization rather than disruption of the axonal cytoskeleton within the AS and support a role of cytoskeletal regulatory proteins in the AS formation.

### Adaptation of the axonal transport in response to injury

If the cytoskeleton is reorganized rather than disorganized immediately following injury then axonal transport and function should not be disrupted. To test this hypothesis, we assessed microtubule dependent transport of two well-established axonal cargoes transported by different molecular motors, APP-GFP or synaptophysin-GFP, and of an organelle, mito-GFP. We acquired movies of axonal transport prior to, during and after axonal injury, and developed a method to track particle movement in bending axons during the injury (Figure 7).

**Figure 7.**
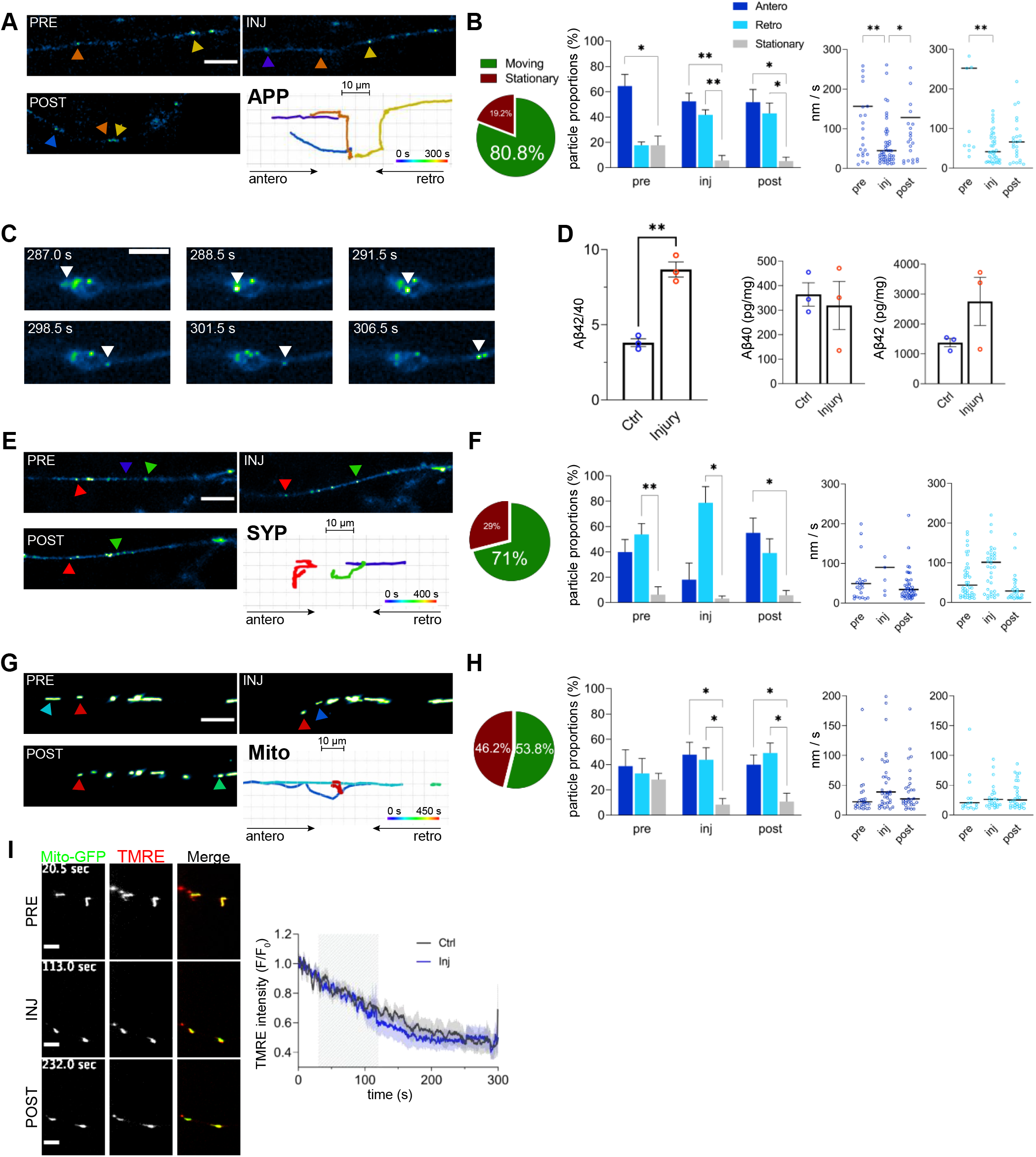
Adaptation of the axonal transport in response to the injury. (A and B) Tracking of APP-GFP particles in pre, during and post-injury stages (A, scale bar: 10 µm). Proportion of moving and stationary particles in the full tracking time (pre, injury and post injury stages) (B). Anterograde, retrograde and stationary movement particle proportions for each stage (n=4). Anterograde and retrograde median velocities in the three different stages (n=33-56). (C) Sequence of an APP-GFP particle moving anterogradely through an axonal swelling (scale bar: 5 µm). (D) ELISA Aβ40 and Aβ42 quantification in the axonal compartment comparing control and injured axons (n=3). (E and F) Tracking of Synaptophysin-GFP particles in pre, during and post-injury stages (E, scale bar: 10 µm). Proportion of moving and stationary particles in the full tracking time (F). Anterograde, retrograde and stationary movement particle proportions for each stage (n=4). Anterograde and retrograde median velocities in the three different stages (n=13-40). (G and H) Tracking of Mito-GFP labeled mitochondria in pre, during and post-injury stages (G, scale bar: 10 µm). Proportion of moving and stationary particles in the full tracking time (H). Anterograde, retrograde and stationary movement particle proportions for each stage (n=5). Anterograde and retrograde median velocities in the three different stages (n=21-40).

APP-GFP particles demonstrated increased retrograde axonal transport at the expense of reduced number of stationary particles during and following injury. This was accompanied by decreased anterograde and retrograde average velocities during injury (Figures 7A and 7B; Video S8). Axonal transport of the APP particles occurred without interruptions also within AS forming during injury (Figure 7C and Video S9). Since post-mortem work indicated perturbed proteolytic processing of the APP during axonal injury (Chen et al., 2004), we next tested APP processing and found significantly increased Aβ42/40 ratio in the axonal fraction (Figure 7D). Conversely, synaptophysin particles showed no significant changes in the directionality during injury, however, they did exhibit significant increase in anterograde and decreased retrograde movement following injury (Figures 7E and 7F; Video S10). This was not accompanied by changes in velocities.

We last examined axonal transport of mito-GFP including mitochondrial labelling with a membrane potential sensor, the tetramethylrhodamine ethyl ester perchlorate (TMRE). During and following injury, we recorded significant increase of both anterograde and retrograde mitochondrial transport, while no changes were detected in average velocities (Figures 7G and H; Video S11). Mitochondrial membrane potentials also showed no differences in the normalized TMRE intensities of mitochondria prior to, during and following axonal injury (Figure 7I; Video S12). These functional assessments of the axons indicate that axonal transport and function do not undergo any major impairments during and following axonal injury.

## DISCUSSION

This study dissects events and mechanisms during the earliest axonal response to physical injury. We found a rapid appearance of AS as a result of the mechanical stress. AS were characterized by an increased spacing of the actin-spectrin rings and a conformational reorganization of the microtubule and neurofilaments tracks, concomitant to phosphorylation changes of cytoskeletal regulators. AS became stable and endure in time only if sustained by high axonal Ca^2+^ concentrations. Finally, we observed that axonal transport and function are preserved throughout and following injury, suggesting that initially AS develop to safeguard axonal homeostasis.

### Axonal injury system

Our model has unique features including: (i) exclusively axonal changes can be followed during and after injury in real-time, (ii) axonal material subjected to injury can be isolated for biochemical, transcriptomic or proteomic studies, and (iii) the use of human derived mature and functional neurons makes the system a better option for clinically relevant studies of axonal injury. In addition, our system has properties that make it comparable to other models, such as the applied force and the size of the AS. During injury, we generated a maximum stress of 40 Pa, which is in the range of other *in vitro* axonal injury systems(Kilinc et al., 2008; Gu et al., 2017) and the sizes of AS that we report both by axolemmal staining and by SEM are consistent with previous *in vivo* studies (Greer et al., 2013).

### Mechanism of AS formation

In light of our experiments, we can distinguish two consecutive stages of the generation of AS. A first stage, characterized by the initial formation of AS and a second stage that involves the sustenance of the AS following injury. Both of these stages are normally accompanied by increased axonal Ca^2+^ levels, which is in agreement with previous reports (Stirling et al., 2014; Staal et al., 2010). However, our pharmacological experiments demonstrated that the role of Ca^2+^ becomes critical for the perpetuation of the AS, while the formation of AS during the injury does not depend entirely on Ca^2+^. Initial formation of AS could be explained as the consequence of fast and reversible mechanisms, that can be the result of either osmotic changes inside the axon (Pullarkat et al., 2006) or of a contractility or relaxation mechanism derived from MYH9 or MYH10 interaction with the axonal cytoskeleton (Costa et al., 2020; Fan et al., 2017). In addition, we identified phosphorylation changes of formins (FMN2), myosin (MYH10) and predicted an upregulation of the Rho/ROCK pathway. These are previously described components of the contractile actin cortex which participates in plasma membrane blebbing mechanism (Charras et al., 2006; Hannemann et al., 2008), and can explain a relatively fast, reversible and Ca^2+^ independent AS formation.

The requirement of high Ca^2+^ levels to sustain AS in time suggests that Ca^2+^ underlies stable changes like the rearrangement of the axonal cytoskeletal components, which may not be promptly reversible. Phosphoproteomics predicted an upregulation of Ca^2+^ dependent kinases immediately following injury that were previously reported to regulate cytoskeletal dynamics, namely PRKCA/B (Larsson, 2006), CAMK1 (Wayman et al., 2004), 2A and 4. Besides, we observed upregulation in JNK1, CDK5, GSK3A/B and ROCK1/2, all which have been previously described to participate in axonal homeostasis (Tararuk et al., 2006) (Reinhardt et al., 2019; Amano et al., 2010). Indeed, through phosphoproteomics we found that the main groups of proteins subjected to regulation via phosphorylation after injury are components of the cytoskeleton (e.g. ANK2, ANK3, ADD3, NEFM, SPTBN1, SPTAN1) or participate in cytoskeleton regulation (e.g. DMTN, SVIL, FMN2, STMN3, DIP2B). Our analysis of cytoskeletal changes shows that the spacing of actin-spectrin rings is increased in the AS, but not in the adjacent shaft, which suggests that actin and spectrin rings participate in the enlargement of the AS membrane. It has been recently shown that actin and myosin work together to maintain the axonal diameter (Costa et al., 2020; Fan et al., 2017), and that α-adducin cap regulates the diameter of the axons (Leite et al., 2016). In line with these studies, our experiments found phosphorylation changes in several actin-associated proteins, including ADD3 and MYH10. Moreover, microtubules and neurofilaments showed no signs of disintegration in AS, but a clear spatial reorganization supporting the transport of cargoes along the axon. Several proteins that participate in microtubule binding (eg. STMN3, MAP1B, DYPSL2, DIP2B, BRSK1 or CAMSAP) were also found to be differentially phosphorylated, strongly suggesting an active process of cytoskeletal remodeling as a result of the injury.

After injury we observed a periodic distribution of the AS along the shaft. This suggests that proteins along the axon involved in some particular process determine the location where AS develop. Previous studies have reported specific Ca^2+^ increases in regions of the axons where AS develop. These regions were characterized by higher abundance of NCX1 and N-type VGCC (Barsukova et al., 2012) or TRPV4 (Gu et al., 2017) channels, which allow Ca^2+^ entry under appropriate conditions. In contrast, our results show that focal elevated Ca^2+^ levels do not necessarily determine the presence of AS. To explain the location where the AS develops, we found a candidate in CAMSAP. This protein presents a stretch-like disposition along the axons, exhibits microtubule minus-end stabilizing properties and also binds spectrin as well as calmodulin. By immunostaining, we observed that CAMSAP2 presents a periodic distribution after injury, reminiscent to the swelling distribution. Furthermore, it was usually present in the AS following a snake-like disposition and interestingly, phosphorylation changes in CAMSAP were found also after injury. Further studies are needed to understand whether CAMSAP determines the location of the AS in the axonal shaft through microtubule and/or cortical cytoskeleton changes.

### AS and pathology

The present work focused largely on the real-time examination of the immediate response of the axons to the injury and showed that in the earliest stages of the axonal injury, AS represent an adaptation rather than an impairment of their structure and function. This concept is also postulated in recent studies where it was shown that in Purkinje cells AS develop as a physiological response in normal development, or compensatory response in disease (Lang-Ouellette et al., 2021; Babij et al., 2013). In addition, some effects that resulted from injury in other studies, were not present in our model. Among them, mitochondrial membrane potential remains unchanged during the injury and mitochondrial transport is not disrupted. There was a lack of significant membrane mechanoporation (Kilinc et al., 2008), and we didn’t observe an intra-axonal increase in Na^+^ or K^+^ levels during injury (von Reyn et al., 2012). Finally, regulation of kinases or proteins involved in signaling of cell death or neurodegenerative pathways was not present.

It’s possible that any of the cytoskeletal, phosphorylation or transport changes that we observed are the starting point that can evolve and acquire pathological characteristics. Even though the cytoskeletal rearrangements are functional in terms of axonal transport, they can be deleterious to the axonal homeostasis if they are not reverted in the long run. For example, we didn’t observe APP accumulation during or immediately following injury, but we observed an increase in the intra-axonal Aβ42/Aβ40 ratio. Given the complex relationship between transport and APP processing, this could be the first step to aberrant Aβ accumulation as observed in a stretch *in vitro* model or in the post-mortem AS of individuals with TBI (Chaves et al., 2021; Johnson et al., 2013). Also, immediately after injury, we observed phosphorylation changes in MAP1B (S1396 and S1400) and in NEFM (S837) in the axonal fraction. Interestingly, these sites were previously reported to be specifically regulated in phosphoproteomics analysis of Alzheimer’s disease brains (Rudrabhatla et al., 2010; Rudrabhatla et al., 2011). On the long run, these phosphorylation changes in fundamental proteins of the axonal cytoskeleton could result in impaired axonal homeostasis. Exploration of this mechanism may lead to understanding of the contribution of TBI as a risk factor for later development of AD (Plassman et al., 2000). Last but not least, it can well be that repetitive or chronic injury is required to produce pathological AS. If this is the case, all the observed changes in cytoskeleton and transport should, through time, be reverted to normal.

Although further studies are needed to decipher the mechanisms of transition between physiology and pathology of the AS, the observed structural reorganization with preserved function of AS following injury offers a window of opportunity for prevention and treatment of brain trauma and other axonal pathologies. Altogether, our findings lay down a molecular, structural and functional framework necessary to build a comprehensive mechanistic understanding of the axonal response to injury from the earliest moments to the post-injury period.

## ACKNOWLEDGEMENTS

We thank members of the Stokin Lab for support and feedback. We thank K. Texlova, A. Wilson and V. Mask for technical contribution. We thank Dr. A. Pompeiano and Dr. S. Katina for support in analysis, and Dr. A. Rainer for computational modeling assistance. We thank Dr. C. Hoogenraad for providing CAMSAP2-GFP construct. We thank Dr. M. Mistrik and Dr. M. Hajduch of the Institute of Molecular and Translational Medicine of Palacky University for Superresolution Microscopy support. We thank D. Gütl and the Electron Microscopy Facility of the Scientific Service Units of IST Austria. We thank Oumayma Essid and PORSOLT for swellings detection algorithm development and analysis. This study was supported by the European Regional Development Funds No. CZ.02.1.01/0.0/0.0/16_019/0000868 ENOCH grant and European Regional Development Fund-Project MAGNET (Number CZ.02.1.01/0.0/0.0/15_003/0000492).

## AUTHOR CONTRIBUTIONS

Conceptualization, V.M.P.D. and G.B.S.; Methodology, V.M.P.D., V.L. and G.B.S.; Formal Analysis, V.M.P.D., J.C. and M.B.L.; Investigation, V.M.P.D., V.L., M.F., P.B., M.Č., I.G.O, N.D., and M.S.; Data Curation, V.M.P.D. and G.B.S.; Writing, V.M.P.D. and G.B.S.; Supervision, G.B.S.; Project Administration, G.B.S.; Funding Acquisition, G.B.S.

## DECLARATION OF INTERESTS

V.M.P.D., V.L. and G.B.S. declare that part of the present work was filed as international patent application: PCT/CZ2019/050063 Pozo Devoto V.M., Lacovich V. and Stokin G.B. (2019): System and method for axonal injury assays.

### Data and materials availability

All data are available in the supplement; raw data are available through the Sequence Read Archive, accession number xxx. Analyzed data and visualizations are available at https://.

## SUPPLEMENTARY FIGURE LEGENDS

**Figure S1.**
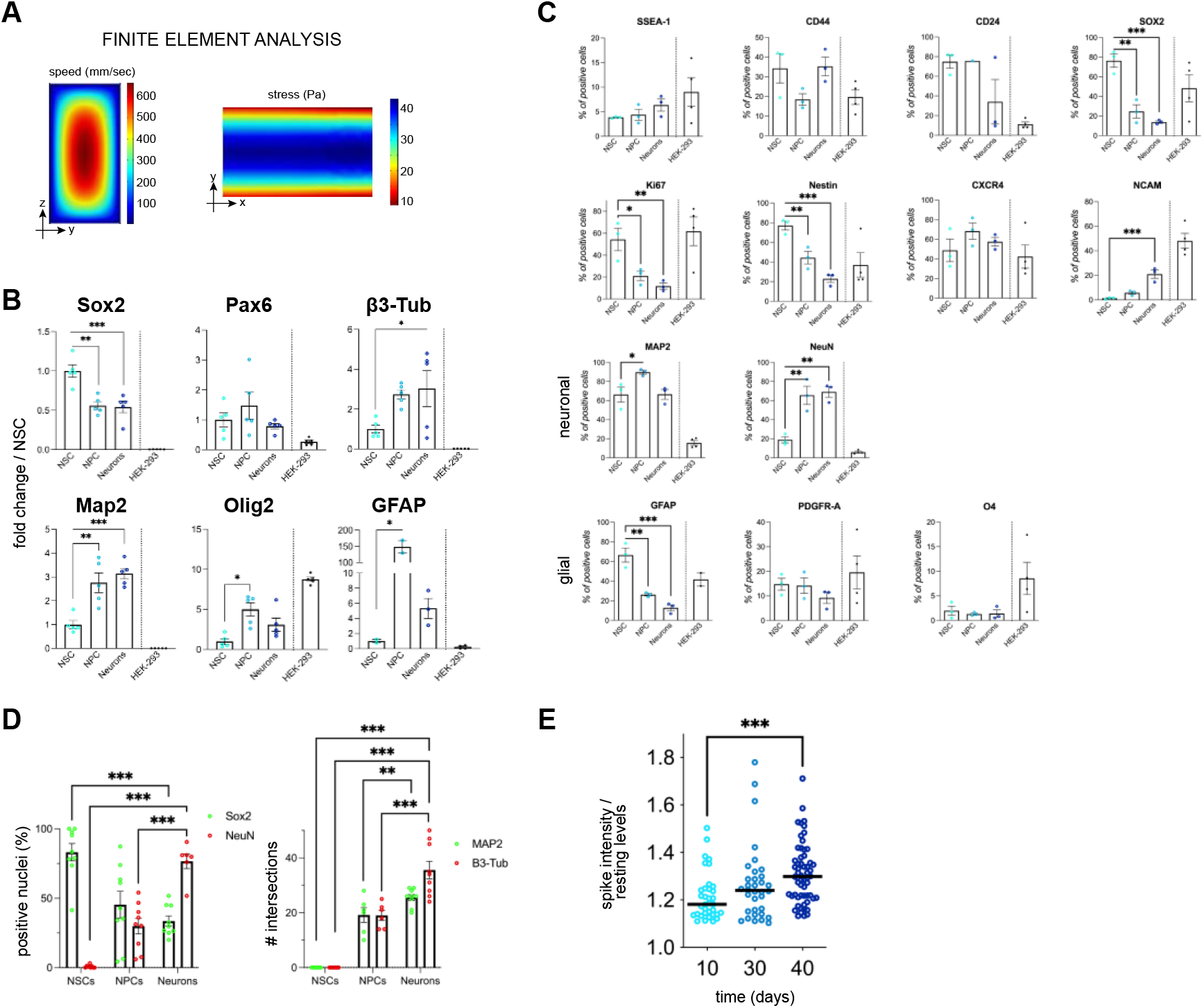
Characterization of injury system, Related to Figure 1. (A) Finite element analysis method applied to a simplified version of the microfluidic chamber used to predict the speed and stress on the injury channel. (B) RT-PCR analysis of mRNA expression levels of markers of early stages, neuronal and glial lineage through differentiation of NSCs to mature neurons. HEK-293 cells were used as a control cell line (n=5). (C) Quantification of positive cells by flow cytometry for different markers of neuronal lineage through differentiation of NSCs to mature neurons. HEK-293 cells were used as a control cell line (n=3). (D) Quantification of immunofluorescence stainings against Sox2 and NeuN or Map2 and β3Tub in NSC, NPC and neuronal stages (n=2, >6 images/n). Number of intersections found from β3-Tub or Map2 positive projections was evaluated (E) Spike intensityies of Ca^2+^ transients (n>34 axons). Data are mean±SEM (B,C and D) or median (E) (*p<0.05, **p<0.01, ***p<0.001). Statistical comparisons were performed using repeated measures one-way ANOVA followed by Dunnett’s multiple comparisons test (B and C), or a one way ANOVA followed by Dunnett’s multiple comparisons test (D), or Kruskal Wallis test followed by a Dunn’s multiple comparison test (E).

**Figure S2.**
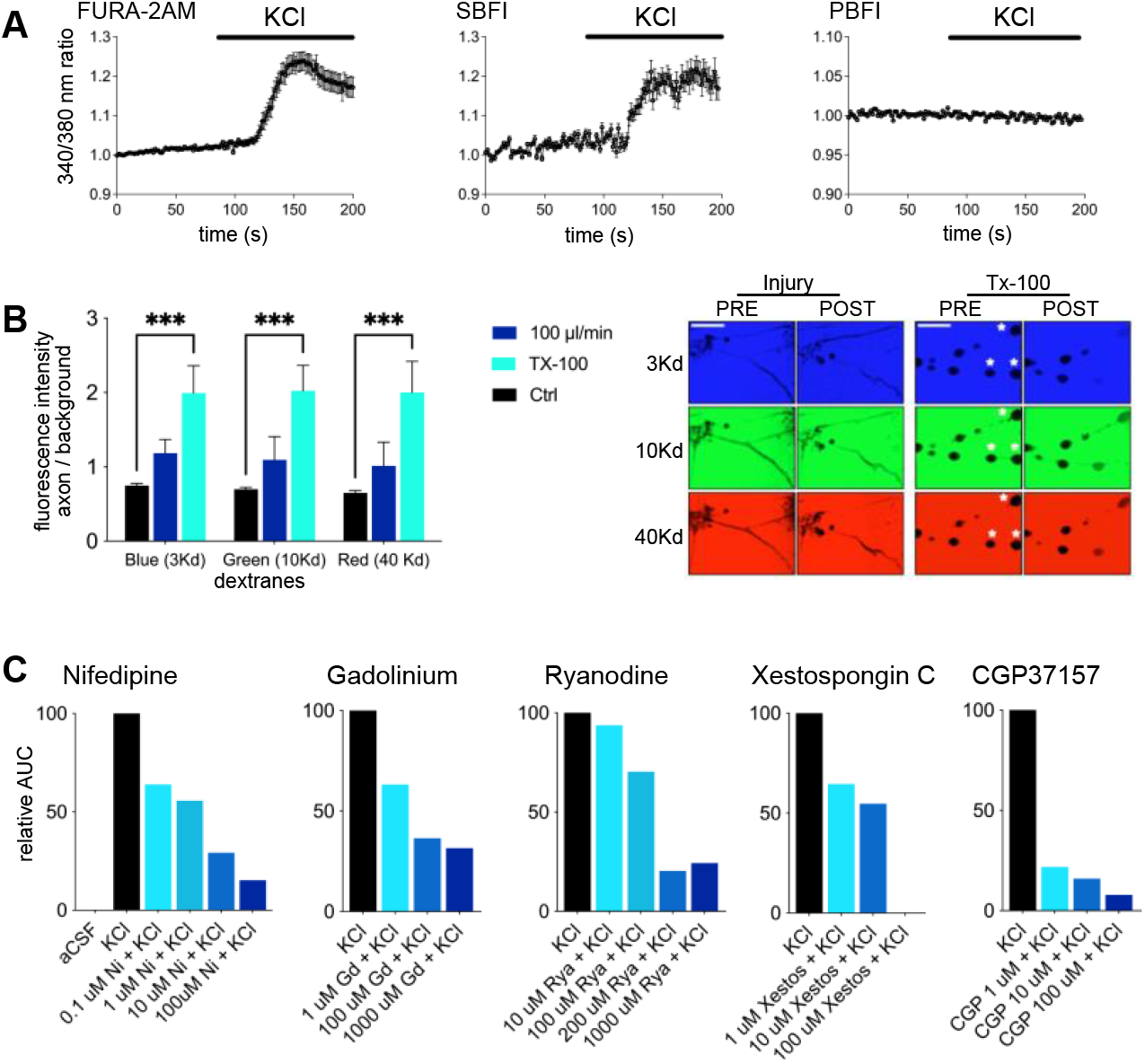
Membrane permeability and Ca^2+^ blockers dose response assessment, Related to Figure 3. (A) Neuronal cultures were perfused with KCl-aCSF to assess the response of the ratiometric probes. PBFI lack of response is due to equilibration of inner and outer K+ concentrations after KCl application (n=15 axons). (B) Permeability of dextranes of different sizes in the axon after injury. Bar graph shows the ratio between the mean fluorescence intensity inside the axon over the mean intensity in adjacent media for each dextrane. Representative micrographs showing the dextranes intensity levels inside and outside the axons, before or after the injury (treatment). Note how parts of the axons (asterisks) get filled with dextranes present in the media(n=3, 7 axons/n, scale bar: 20 µm). (C) Dose response curves of Ca^2+^ blocker compounds in neuronal cultures. Fluo-4AM intensity was measured and area under the curve calculated after application of 50 mM KCl (n=15 axons). Data represented as mean +SEM (***p<0.001). A two way ANOVA was performed followed by a Dunnett’s multiple comparisons test versus the control treatment (B).

**Figure S3.**
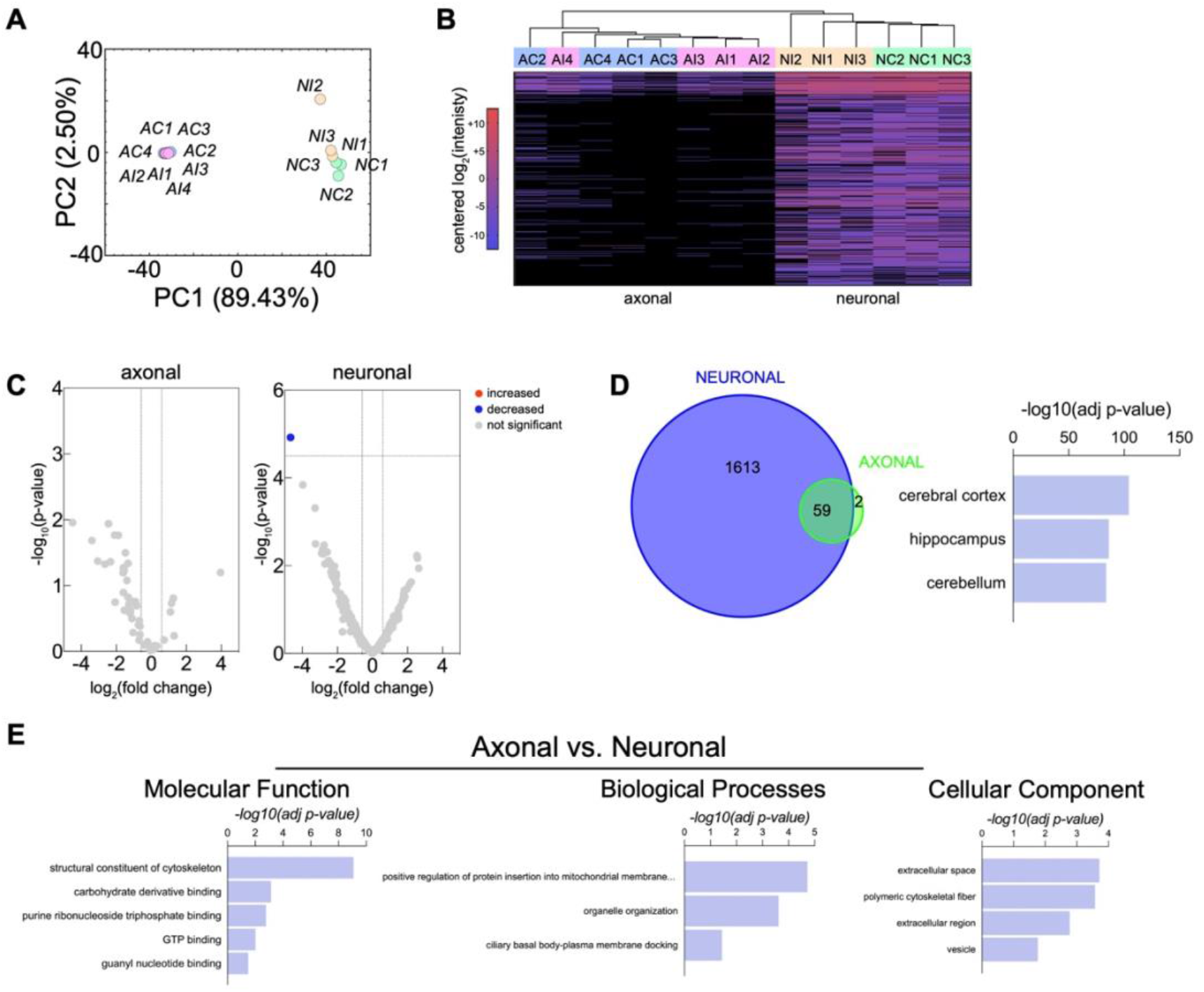
Axonal proteomic profiling before and immediately after injury, Related to Figure 4. (A) PCA of axonal and neuronal fractions (A/N) in both control and injury treatments (C/I). n=4 axonal fractions, n=3 neuronal fractions. (B) Heatmap of differentially expressed proteins. (C) Volcano plots depicting the quantitative changes of injury compared with control treatments for the axonal and the neuronal fractions. (D) Total number of proteins present in the neuronal and axonal fractions (left panel). Human Protein Atlas top terms of the enrichment analysis for the neuronal fraction proteins (right panel). (E) Enrichment analysis of the axonal fraction proteins using as background the neuronal fraction proteins. Significant terms for each GO category.

**Figure S4.**
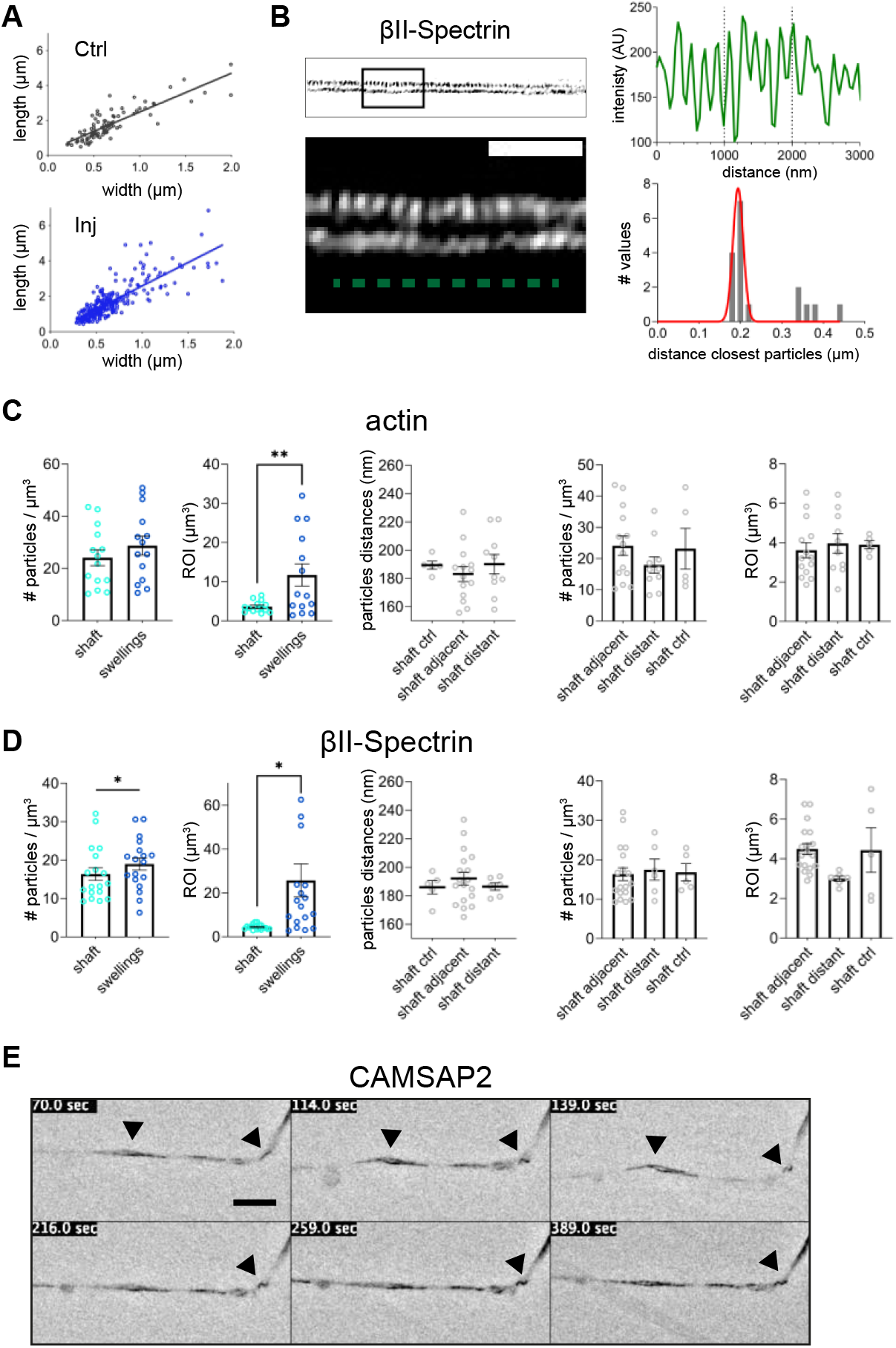
Structural reorganization of the axonal cytoskeleton following injury, Related to Figure 6. (A) Correlation between the width and the length of the AS present in control and injured conditions in scanning microscopy images (control n=96, injury n=260). Linear regression for control and injured, y=2.23x+0.2574, R^2^=0.73 and y=2.63x+0.04, R^2^=0.65. (B) Validation of the closest neighbour distance model to analyze distance between particles. Upper plot shows the profile of intensity through the axon long axis, showing inter-peaks distances of approximately 200 nm. Lower plot shows the closest neighbour distance analysis of the same axon, with a median of 200 nm. Scale Bar: 1 µm. (C and D) Actin (C) and βII-Spectrin (D) particles densities and volumes for the different ROIs analyzed in Fig. 6E. Comparison of particles distances in shafts of axons that were not subjected to injury versus region of the shaft distant from the swelling versus region of the shaft adjacent to the swelling. (E) Movie frames of CAMSAP2 transfected neurons showing the bending (arrowheads) of the CAMSAP2 filaments during and after injury (scale bar: 10 µm). Data are mean±SEM (*p<0.05, **p<0.01). Statistical comparisons were performed using the student t-test and one way ANOVA.

